# Psychophysical evidence and perceptual observations show that object recognition is not hierarchical but is a parallel, simultaneous, egalitarian, non-computational system

**DOI:** 10.1101/2021.06.10.447325

**Authors:** Moshe Gur

## Abstract

Object recognition models have at their core similar essential characteristics: feature extraction and hierarchical convergence leading to a code that is unique to each object and immune to variations in the object appearance. To compare computational, biologically-feasible models to human performance, subjects viewed objects displayed at a wide range of orientations and sizes, and were able to recognize them almost perfectly. These empirical results, together with consideration of thought experiments and analysis of everyday perceptual performance, lead to a conclusion that biologically-plausible object perception models do not even come close to matching our perceptual abilities. We can categorize many thousands of objects, discriminate between enormous numbers of different exemplars within each category, and recognize an object as unique although it may appear in countless variations―most of which have never been seen. This seemingly technical, quantitative failure stems from a fundamental property of our perception: the ability to perceive spatial information instantaneously and in parallel, retain details including their relative properties, and yet be able to integrate details into a meaningful percept such as an object. I present an alternative view of object perception whereby objects are represented by responses in primary visual cortex (V1) which is the only cortical area responding to small spatial elements. The rest of the visual cortex is dedicated to scene understanding and interpretation such as constructing 3D percepts from 2D inputs, coding motion, categorization and memories. Since our perception abilities cannot be explained by convergence to “object cells” or by interactions implemented by axonal transmissions, a parallel-to-parallel field-like process is suggested. In this view, spatial information is not modified by multiple neural interactions but is retained by affecting changes in a “neural field” which preserves the identity of individual elements while enabling a new holistic percept when these elements change.

## Introduction

We can easily recognize many thousands of objects even though each may appear at any one of a huge number of variations. This ability, which is essential for normal behavior, has been intensely studied and a plethora of possible object recognition mechanisms has been offered. Many such mechanisms are either inspired by the visual system or aim to emulate its anatomy and physiology (see reviews by DiCarlo et al., 2012; Rolls, 2012). It is easy to see that almost all models do not strictly follow the functional anatomy of the visual cortex but, at best, are only inspired by some physiological or perceptual principles. For example, the very popular and very successful deep-learning approach (cf. LeCun et al., 2015) with its many layers, reliance on a huge number of exemplars, and massive connectivity coupled with guided “learning”, has little in common with the actual makeup of the brain. Other models such as the recognition by components (Biederman, 1987) are inspired by psychophysical data but are not biologically plausible. Moreover, practically all models have at their core similar essential characteristics: feature extraction and hierarchical convergence leading to a code that is unique to each object and immune to variations in the object appearance. Since I claim that these characteristics are not compatible with present results as well as with fundamental perceptual observations, it is sufficient to consider the most influential, biologically-based model: sparse population coding by expert cell ensembles (ECE).

### Sparse population coding by expert cell ensembles (ECE)

This dominant physiology-based view of object representation and recognition posits that, first, the image is analyzed into its basic elements, such as edges or line segments, by feature-selective cells in the primary visual cortex (V1). Then, after several steps of hierarchical convergence and integration of simple elements, small ensembles of expert cells, by their collective responses, represent objects uniquely and invariantly. Recognition is achieved, presumably, by comparing the ensemble’s acute firing pattern to a memory one.

Hubel and Wiesel’s (1962, 1968) evidence of orientation selective cells in the cat and monkey V1 established the basic tenets of the ECE hypothesis. They suggested that feature selectivity in V1 cells is achieved by hierarchical convergence of cells with concentric receptive fields (RFs) that generates “simple” cells with elongated RFs which, in turn, converge to generate the next hierarchical link―”complex” cells. They predicted that further convergence in areas downstream from V1 will enable encoding of increasingly complex features and at the same time allow a considerable degree of invariance.

The next 2 areas downstream from the monkey V1, V2 and V4, show only a modest increase in feature selectivity (Anzai, 2007; Hegde and Van Essen, 2007; Gallant et al., 1996; Carlson et al., 2011; Roe et al., 2014), and it is the temporal cortex where cells that clearly respond to elaborate integrative features are found. Numerous studies showed that cells in the monkey inferotemoral (IT) cortex are selective for various complex shapes including faces (Perrett et al., 1982; Orban, 2008; Freirwald and Tsao, 2010; Eifuku et al., 2011; Hung et al., 2005; Yamins er al., 2014; Ohayon et al., 2012; Sato et al., 2013; Hirabayashi et al., 2013). The hierarchical transformations leading to shape-selective cells are paralleled by an increase in spatial integration from cells integrating over a few minutes of arc in V1 to cells at the pinnacle of the hierarchy responding within a large portion of the visual field (VF). It is thought that the collective properties of “expert” cells enable individual objects to be recognized in spite of changes in global parameters such as size or viewpoint (Freiwald and Tsao, 2010), although, it should be noted, only “face-ensembles” were found but not ensembles specializing in other visual objects. Research in homologous areas in human visual cortex is consistent with single cell data from the monkey IT cortex (Axelrod and Yovel, 2012; Parvizi et al., 2012; Anzellotti et al., 2014; Ramirez, 2014). Experimentalists invariably identify the temporal cortex as the site for object representation and recognition (Freirwald and Tsao, 2010; Eifuku et al., 2011; Hung et al., 2005; Sato et al., 2013; Hirabayashi et al., 2013; Axelrod and Yovel, 2012; Parvizi et al., 2012).

Fundamental to the current view of object representation are two main processes—encoding and hierarchical convergence. It assumes that once parallel retinal spatial information reaches feature selective V1 cells, it is transformed into a code carried by cells that by hierarchical convergence respond to increasingly elaborate features. Suggested codes may be relatively simple whereby single cells’ responses encode spatial features such as lines or faces, or more complex ones where patterns of responses by ensemble cells encode the required spatial information. How such a code is decoded into our detailed, parallel space perception is usually not dealt with.

Note that since according to the ECE hypothesis objects are represented by the firing patterns of corresponding ensembles, object discrimination, categorization, and recognition must all rely on these patterns.

In summary, the main tenants of the ECE hypothesis are:

1) ***Category-specific ensembles***. Objects are represented by expert-cell ensembles where each ensemble responds vigorously only to a single object category (e.g., human faces), 2) ***Category uniqueness and invariance***. These category-specific responses are immune to changes in objects appearance such as size, orientation, or brightness, 3) ***Individual uniqueness and invariance***. Each individual member within a given category generates a response that differs from responses to any other category member and is immune to changes in its appearance.

To test the plausibility of the ECE hypothesis, I studied how well subjects can recognize briefly presented objects belonging to three classes (doodles, animal contours, and faces) shown at many sizes and orientations. The empirical results and a review of perceptual observations unequivocally show that the ECE hypothesis as well as any computable mechanism relying on convergence, feature extraction, and object-specific codes, is not biologically feasible and that a different, parallel, non-computable mechanism must be considered.

## Methods

### Stimuli

There were three stimulus categories: meaningless doodles, animal dot-contours, and black & white face photographs (Fig. 1). Each category consisted of four different exemplars. Each exemplar was presented at 12 orientations (0, 30 … 330º), and 10 widths (0.6, 0.8 … 2.4º). Stimuli were displayed on a 60 Hz, 24” monitor with 1920×1080 pixels. The screen was viewed binocularly with natural pupils and a chin rest from a distance of 114 cm.

**Figure 1.**
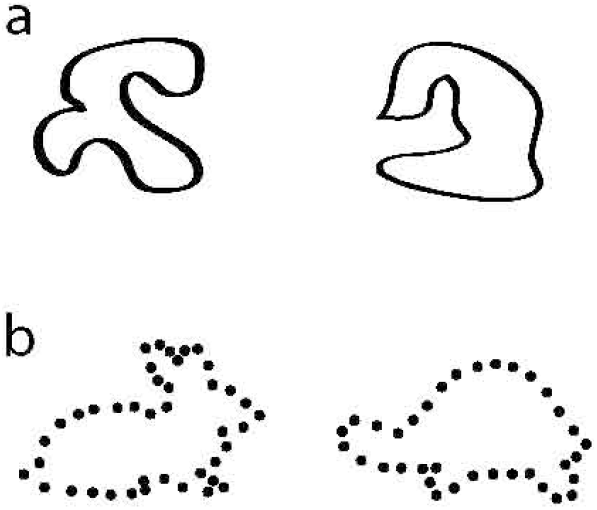
Examples of the doodles (a), and the animal contours (b) stimuli. Examples of the face stimuli can be seen in Gur (2018), Fig. 1.

### Doodles

Hand-drawn shapes were scribbled by the author such that there would not be any clear resemblance to familiar objects or any preferred orientation (Fig. 1a).

### Contours

Outlines of four animals (turtle, rabbit, bird, and shark) were followed using black dots (Fig. 1b).

### Faces

Black & white photographs of four individuals of the approximate same age were used. Great care was taken to ensure that the faces differed only by the characteristics and arrangement of facial features (mainly the eyes, nose and lips) and not by coarse elements such as the differences between short and long head hair, hair style, glasses or facial hair (Fig. 1c; see Gur, 2018 for details). The target faces were displayed within a grey background of the same average luminance as the image (55 cd/m^2^).

### Masks

Masks were constructed from randomly arranged elements taken from each stimulus category: line segments for doodles, dots for contours, and face segments for faces. The masks size (3×3º) was larger than the largest stimulus.

### Procedure

#### Familiarization

Prior to testing, the subjects familiarized themselves with the four different exemplars comprising each category. First, they viewed, at their leisure, the four exemplars with the associated keyboard arrows (left, right, up, down). Then a short simulation of the actual experiment where the four exemplars were presented at two different sizes and four orientations was run, and the subjects had to press the corresponding arrow. It is important to note that the objects presented in this phase were different from those presented at the actual experiments; they had different aspect ratios (e.g., narrower and longer), orientations (e.g., 15º), and sizes (e.g., 2.5º). This was done so that all objects presented in the first experimental session were seen, in their exact form, for the first time.

#### Object recognition

A four-alternative forced-choice procedure was used. Subjects had to select a particular object out of four possible ones. In any one session the subject was presented with one object category (doodles, contours, or faces). In each category there were four different objects and each object could be displayed in one out of twelve orientations and one out of ten sizes for a total of (4×12×10) 480 variations. A given object, size, and orientation was randomly selected. Each subject participated in five sessions/category for a grand total of (480 presentations x 5 sessions x 5 subjects) 4800 determinations/category. Each stimulus was displayed for either ∼67 (doodles or contours) or 83ms (faces) followed immediately by a 100ms mask. Two hundred ms after the subject responded, the next stimulus was presented. The order of image presentation was counterbalanced across days and subjects

#### Subjects

Five subjects (three women; four subjects were 23-35 years old. One subject’s age was 69) with normal or corrected to normal vision participated in all experiments. Four were naÏve about the purpose of the experiment. The fifth was the author. The experimental protocol was approved by the Technion ethics review board. All subjects gave their written informed consent before the experiments.

The subjects were instructed to carefully fixate at the center of the screen for the presentation duration. They learned quickly that due to the very short presentation times, only a good anticipatory fixation allows the image to be recognized. The subjects were told that they should respond as fast as possible but that their first priority was to correctly identify the object.

#### Data and analysis

Success rates and reaction times as a function of orientation and size were collected. Since the ability to correctly identify an object was based on a binary correct/incorrect response, success rate was calculated, per subject, for a given orientation across all sizes and all within-category objects, or for a given size across all orientations and within-category objects. Thus each size success rate was the ratio of correct responses for 40 (4 objects x 10 sizes) determinations, each orientation success rate was the ratio of correct responses for 48 (4 objects x 12 orientations) determinations. These ratios were calculated for each session so that the over-all success rate for each orientation and size was averaged over 25 (5 sessions x 5 subjects) single session averages.

Statistical significance between averaged success rates and reaction times (RTs) for different orientations and sizes was calculated by a pairwise two-sided t-test (SAS).

## Results

As noted above, in each initial experimental session, the specific stimuli used in all three categories were seen by the subjects for the first time. In each session, for each category, four objects were presented in one of 12 possible orientations and 10 sizes. Each configuration was seen only once. Since there were four objects in each category, chance recognition level was 25%. The first demo in the supplemental file (Demos.docx) can give the reader an impression of the required task.

### Doodles

In this experiment the subjects viewed four different scribbles which had no clear relations to a concrete object and no preferred orientations (Fig. 1a). As can be seen (Fig. 2) recognition for all orientations was near-perfect (91-96 %; Fig. 2a) and RTs were within a narrow 640-690ms range (Fig. 1b). Size, as expected, had some effects (Fig. 2c, d); in the two smallest sizes recognition was below 90% (87, 89 % for the 0.6, 0.8º stimuli, respectively) and RTs very gradually decreased as stimuli size increased. However, these effects were rather minor. Recognition for the smallest two sizes (combined) were significantly (p<0.02) lower than the two largest ones but not for the first five sizes vs. the five largest ones. There were no significant differences in RTs between sizes.

**Figure 2.**
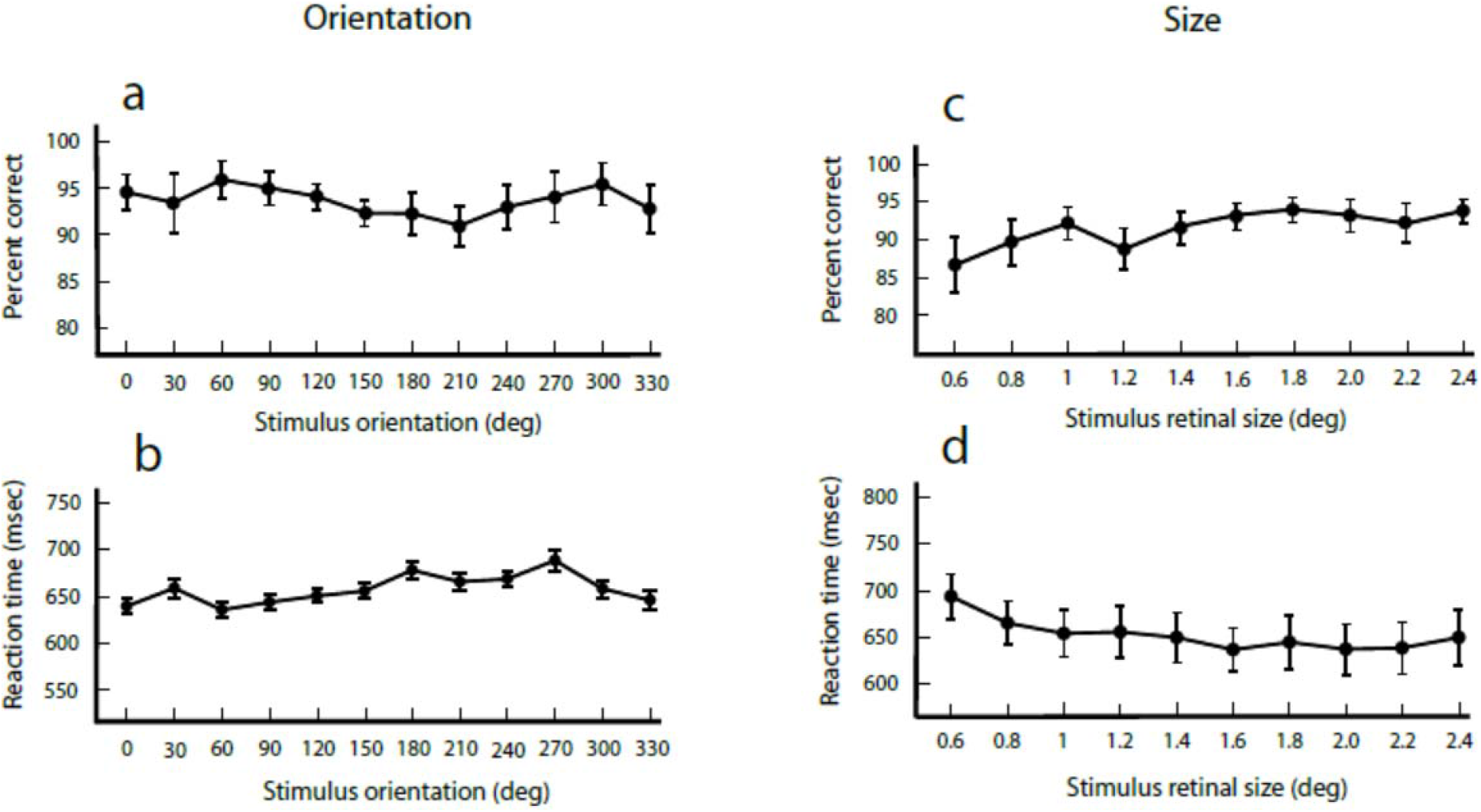
Percentage of correct responses and RTs, to doodles presented at 12 different orientations (a, b, respectively), and 10 sizes (c, d, respectively). Error bars denote the STD.

### Contours

Here, objects were sparsely represented; only contours were displayed and each contour contained ∼ 38 dots (Fig.1b). Yet, recognition rates were very high―all orientations except one were recognized at an 89-95% level with no obvious preference for any given orientation (Fig. 3a). RTs to the various orientations fluctuated within a narrow range (660-710ms; Fig.3b). Although we might have expected that upright orientations would be easier to recognize, both recognition rates and RTs showed no strong preference for the upright position; the lowest recognition rates were at 210 and 240º, and the highest two at 270 and 300º. The shortest RTs were found away from the upright orientation at 30, 60, and 270º while the longest RTs were in response to the 150 and 300º orientations. The differences between recognition rates and RTs were not statistically significant. Size, as in the doodles case, did affect recognition success (Fig. 3c, d); recognition for the smallest two sizes (combined) was significantly lower from that of the largest two sizes (p<0.1). Similarly, recognition success for the smallest five sizes was significantly lower than for the largest five (p<0.001). RTs for the various sizes varied within a very narrow range of 677 to 652ms and the RTs differences were not statistically significant.

**Figure 3.**
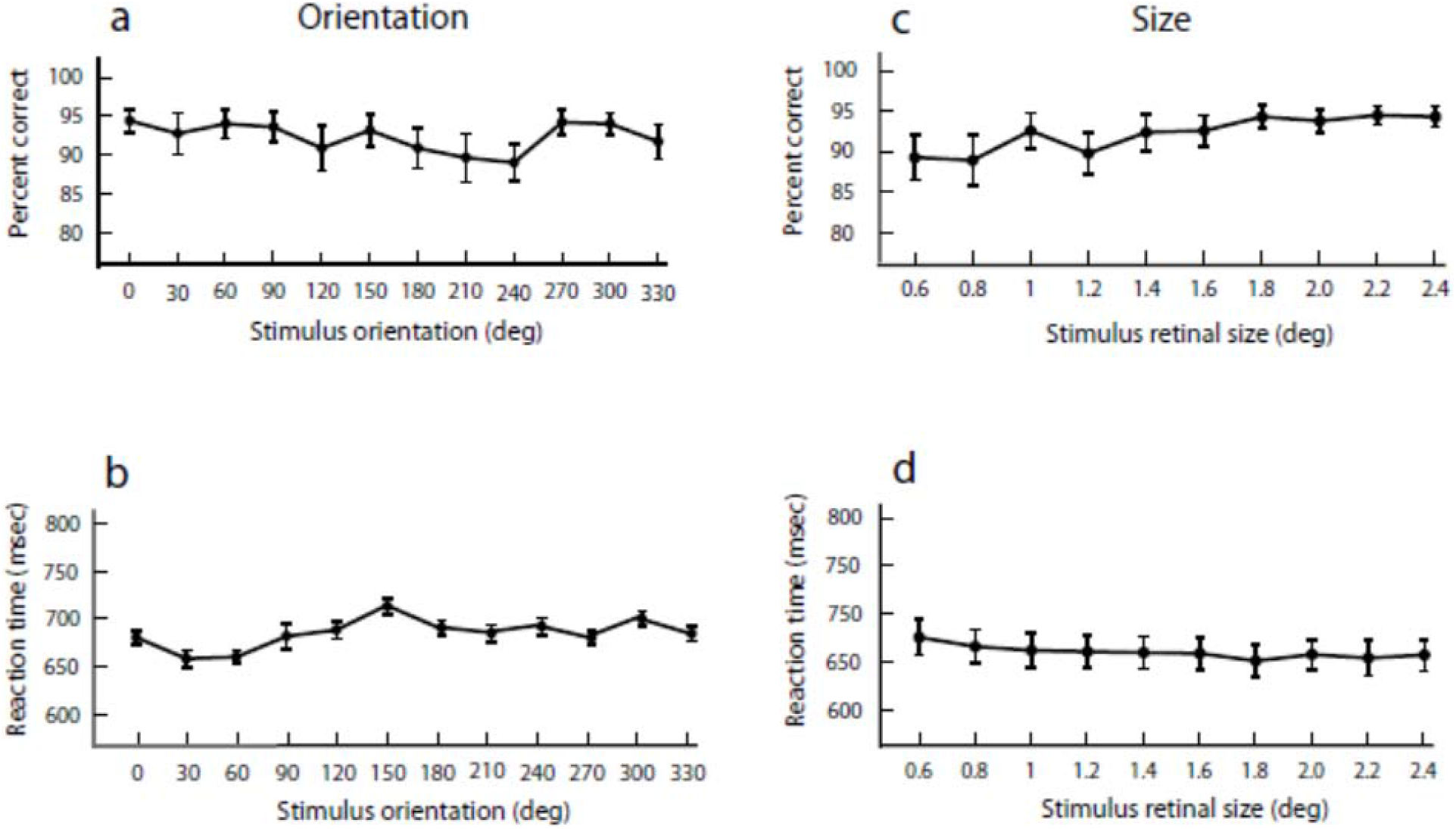
Percentage of correct responses and RTs, to animal contours presented at 12 different orientations (a, b, respectively) and at 10 sizes (c, d, respectively).

### Faces

Recognizing faces presented at various orientations and sizes was a *priory* more challenging than recognizing doodles and contours. The various faces presented here lacked any telltale features such as facial hair or glasses and could only be recognized by the overall arrangements of facial features―the shape and relative positions of the eyes, nose, and lips. Also, it is well known that face perception is impaired as faces get close to the upside-down orientation where it becomes more difficult to detect changes in local features (Yin, 1969). Present data (Fig. 4) do show that face orientation affected recognition; as the orientation of the presented faces got closer to the inverted one, recognition rates dipped and RTs increased. However, recognition of inverted and near-inverted faces was still excellent (87-90% success) and only the lowest recognition rate (87% at 210º orientation) was significantly different (p<0.05) from the highest recognition rates (95%). The more pronounced effect was on RTs that increased by ∼100ms for inverted faces compared to upright ones (Fig. 4b). The three slowest RTs (at 150, 180, and 210º) were significantly slower (p<0.01) than RTs for orientations around the upright position (0, 30, 60, and 330º). Changing face size affected recognition at the smallest sizes (0.6, 0.8º) where the recognition rate was 0.87, and 0.9, respectively―significantly (p<0.01) lower than the average correct responses (0.96) for the four largest faces (Fig. 4c). Although RTs varied over a narrow range (660-690ms) the slowest RTs, although not significantly different from other RTS, were found in responses to the two smallest sizes (Fig. 4d). Such an affect is expected since for the smallest face width (36’), where facial features size is <10’, and individual variations in shape and position are at the 1-3’ range, recognizing a face presented for a mere 83ms is quite a challenge. Nonetheless, performance even for the smallest size was excellent; recognition rate was 0.89 and RT increased by less than 50ms compared with the averaged RTs for the largest two sizes.

**Figure 4.**
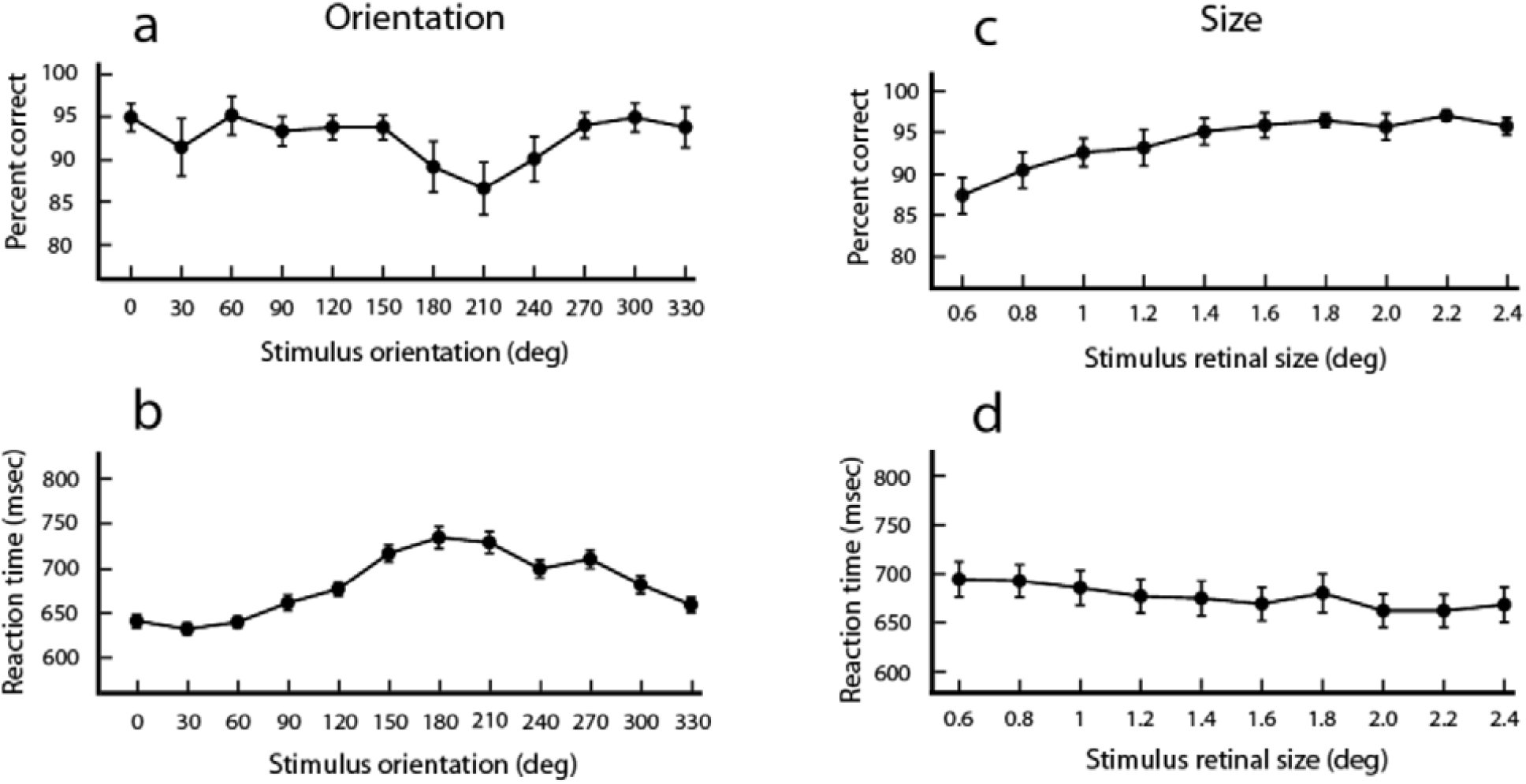
Percentage of correct responses and RTs, to faces presented at 12 different orientations (a, b, respectively) and at 10 sizes (c, d, respectively).

## Discussion

In the present experiments I studied how object recognition is affected by large variations in orientation and size. Three types of objects were used: doodles, animal contours, and faces. All objects were seen for the first time in the familiarization phase which lasted, for each category, less than a minute. Moreover, objects seen during familiarization were different in aspect ratio, size, and orientation from objects presented during the experiments. Thus, during a single experimental session, for each object category, each of the four objects was seen only once in a particular size and orientation. In each session 480 different stimuli (4 objects x 12 orientations x 10 sizes) were presented. What is most impressive about present results is the ease of recognition; even though the various objects were seen for a mere 67ms (drawings and contours) or 83ms (faces), followed immediately by a 100ms mask, and were either seen for the first time (first session) or for a few more times out of thousands configurations (in following sessions), recognition was excellent; for all variations, recognition success rate was >87% and for 58/66 configurations (12 orientations + 10 sizes―for each category) success rate was >90%. Given that the subjects were motivated to respond quickly so that occasional pressing of the wrong key was unavoidable, such rates are near perfect. In other words, the subjects had absolutely no problem in recognizing a particular object regardless of orientation and size.

### Orientation

That orientation did not affect recognition in the doodles case is not surprising given the lack of any obvious orientation preference. Also in the animal contour case where some orientations are more prevalent in real life (Fig. 1), using just contours makes the shape very obvious which allows unequivocal recognition regardless of orientation. The really interesting case is that of human faces where it has been shown that inverted faces are harder to recognize than upright ones (Yin, 1969). Here, indeed, orientation did affect recognition ability but only slightly, so that even for orientations around 180º (Fig. 4a), recognition rates were reduced from ∼92-95% around the upright orientation to ∼87-94% around the inverted one. Unlike doodles and contours where RTs were not affected by orientation, the effect of face orientation on performance was substantial. Even though recognition of inverted faces was excellent, RTs increased from ∼620-630ms for orientations near the upright face to ∼730-740ms for inverted faces (Fig. 4b). This RT increase may reflect the difficulty in selecting the correct face when inverted―although the added difficulty did not have a dramatic effect on recognition.

### Size

Image size ranged from 0.6º which stimulates the very center of the fovea to 2.4º that covers all central vision. Here, even though for all sizes recognition was excellent (87-96%; Fig. 4c), recognition for the 0.6 and 0.8º was somewhat lower (87 and 90%, respectively) reflecting the difficulty in discerning characteristic features that for such small over-all sizes can be as small as a few minutes of arc. RTs varied within the rather narrow range of ∼620 – 690ms. It can be seen thus that for the smallest size success rates were lower and RTs were longer than for larger sizes. This result is only to be expected since for a 0.6º stimulus width, constituent elements, particularly regarding faces, are quite small and thus present some perceptual difficulties.

To summarize, objects seen either for the first time or for a few additional times at many orientations and sizes, for a mere 67 or 83ms (followed by a mask), were recognized at ease. Recognition was thus possible regardless of variations in size and orientation, via a strict feed-forward mechanism, and with no memory patterns to compare to.

As noted in the Introduction, the sparse expert cell ensemble (ECE) hypothesis shares its essential features with all computational models, and since it is the most biologically plausible one, it is sufficient to see whether its structure and predictions are compatible with present results.

### Present results challenge the ECE dogma in several ways

#### There can be no ensembles that are tuned to the specific stimuli used here

The very short exposure times (67, 83ms followed by a 100ms mask) preclude learning by adjustment of synaptic weights that can generate new expert cell ensembles. Also, it is doubtful whether there exists in our brain ensembles that are already tuned to the images used in the current experiments since none of the tested objects was ever seen in its exact form prior to the experimental sessions; doodles and sparse-contour animals are not encountered in every day environment and the face stimuli were quite different from real 3D faces and even from the usual photographs of faces. The latter are characterized by many more features than current stimuli which are defined only by the shape and relative position of the eyes, nose and lips.

#### Biologically realistic ensembles cannot be invariant to present stimuli

The range of sizes (0.6-2.4º) and orientations (0-360º) makes it very unlikely that a sparse ensemble can generate very similar responses to such very large variations in global characteristics―yet distinguish between similar but different objects. For example, Fig. 5A shows the hierarchical convergence in the ventral stream which leads to a large increase in RF size from the very small RFs (1-5’ in the center of vision) in V1 to a 2-3 fold increase in each succeeding layer resulting in very large (10-50º) non-retinotopic RFs found in the IT cortex. Now consider an example from the animal contour experiment (Fig. 5B). Subjects had no difficulty recognizing the turtle and the bird contours as different objects (upper row, left and middle columns) while identifying the two contours shown in the lower row (left and middle columns) as a bird. In the right-hand column of panel B, the two stimuli are superimposed to show the degree of visual field overlap which is considerable for the turtle/bird contours and non-existent for the large/small birds. In panel C the combined stimuli are superimposed on the “foveal” representation of the schematic V1 layer. An IT RF (not to scale) is also depicted. The ECE hypothesis predicts that there are IT animal-expert-cell-ensembles that will generate very similar responses to the two bird stimuli―although they differ greatly in size and orientation and, most importantly, do not share any V1 RFs. The same ensembles, according to the ECE hypothesis, will consistently generate different responses to the bird/turtle stimuli although the two stimuli share almost all their V1 RFs. Furthermore, in our experiments, stimulus size varied between 0.6-2.4º which covers only a small fraction of IT RFs (10-50º) and would thus either generate a relatively weak response for the >1º stimuli or no response for the <1º ones (Freiwald and Tsao, 2010). How then can such weak sub-optimal responses be so fine-tuned to give different responses for the two stimuli sharing, practically, the same visual space yet generate identical responses to two stimuli that share no visual space? Given these considerations it is very unlikely that expert cell ensembles can be invariant to very large changes in stimulus size and orientation yet be able to discriminate between stimuli that differ by very few elements. In fact, it is very hard to think of any biologically plausible mechanism that can accomplish such a feat and, indeed, no one has ever been suggested. All proposed models always employ stimuli that are not very different in global features such as size or orientation but are quite different in local characteristics such as hair.

**Figure 5.**
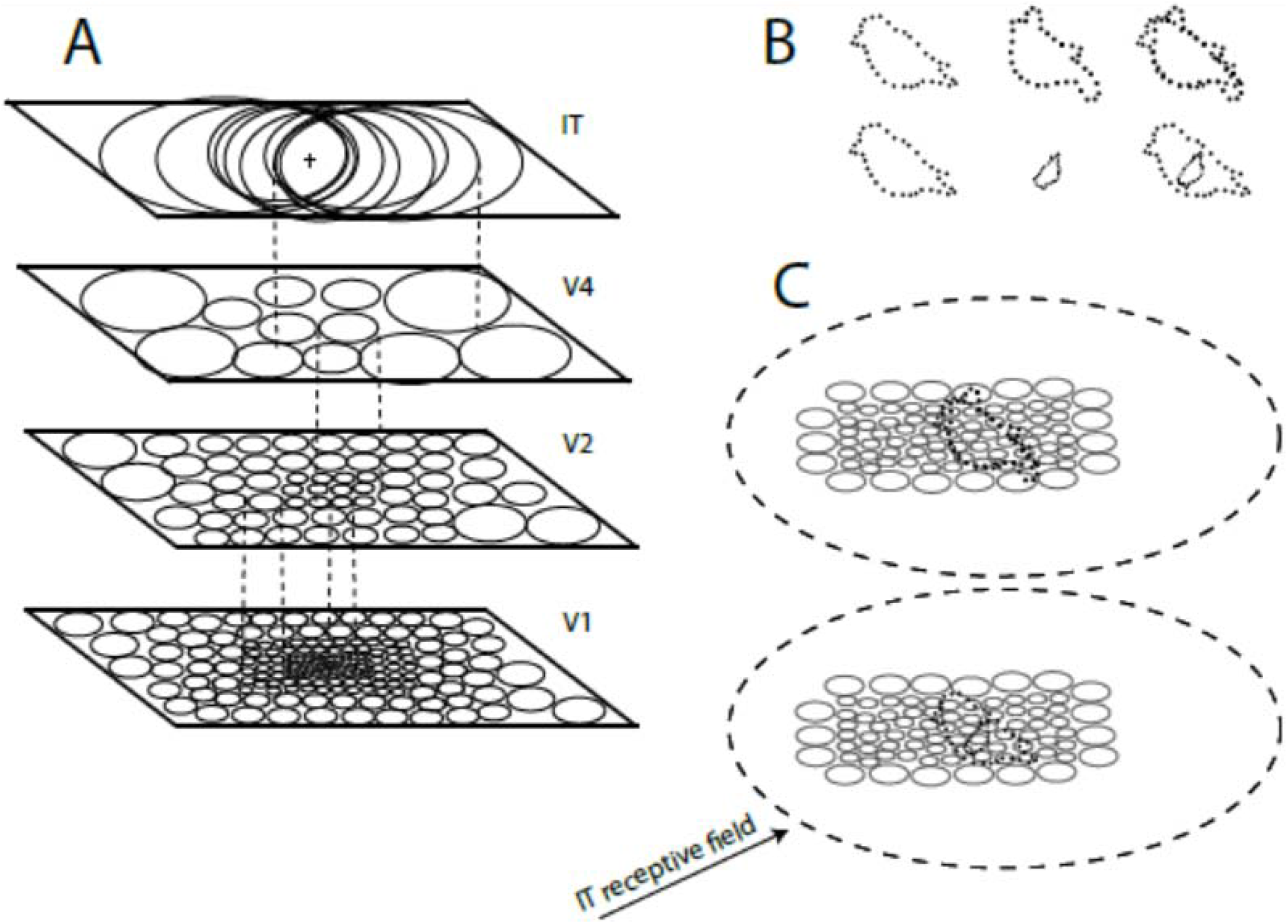
The extent of present stimuli impact within the organization of the visual system. A. Schematic organization of ventral stream areas. “V1” shows RFs within central vision;”V2” shows RFs in central and near periphery visual field (VF);”V4” shows RFs in both central and peripheral VFs, while in “IT”, where retinotopy is lost, RFS encompass large portions of the VF. B. Upper raw: the bird and turtle stimuli shown at equal size, individually, and superimposed. Lower row. Same as the upper one but for different sizes such that in the superimposed case there is no overlap between images. C. The fused images are superimposed on the foveal representation in “V1”. An IT RF is also shown. Although smaller than actual size, it is much larger than V1 foveal representation.

### Going beyond present results ―are the basic tenants of the ECE hypothesis plausible?

#### Can objects be represented by category-specific ensembles?

To show how unlikely is the category-specific premise of the ECE hypothesis, let us consider a simple thought experiment (Fig. 6) where the subject is instructed to ignore constituent elements and consider the over-all shape when identifying an object category. Simple and extremely easy but with fundamental implications. First, there are thousands object categories that could have been presented to the subject: geometric structures, trees, flowers, cars, bicycles, buildings, animals, faces, body parts, furniture, tools, jewelry, etc. According to the ECE hypothesis each object category such as faces is represented and recognized by a specific expert ensemble. Are we then to assume that there are many thousands of distinct ensembles in the IT cortex, each representing a particular object category such as houses or furniture? Incidentally, are chairs represented by a different ensemble than stools, recliners, tables, sofas, sideboards, benches, beds, etc.? Are four-legged chairs represented by a different ensemble than three-legged, one-legged, no legged, egg-shell, stiff-back, or straw-back chairs? It seems thus that for expert-cell ensembles to represent all object categories, thousands, perhaps tens of thousands, are required. Given such an extraordinary biological requirement and the fact that except for “face-ensembles” no other object ensembles were found in the IT cortex, the very existence of “object-ensembles” is highly doubtful.

**Figure 6.**
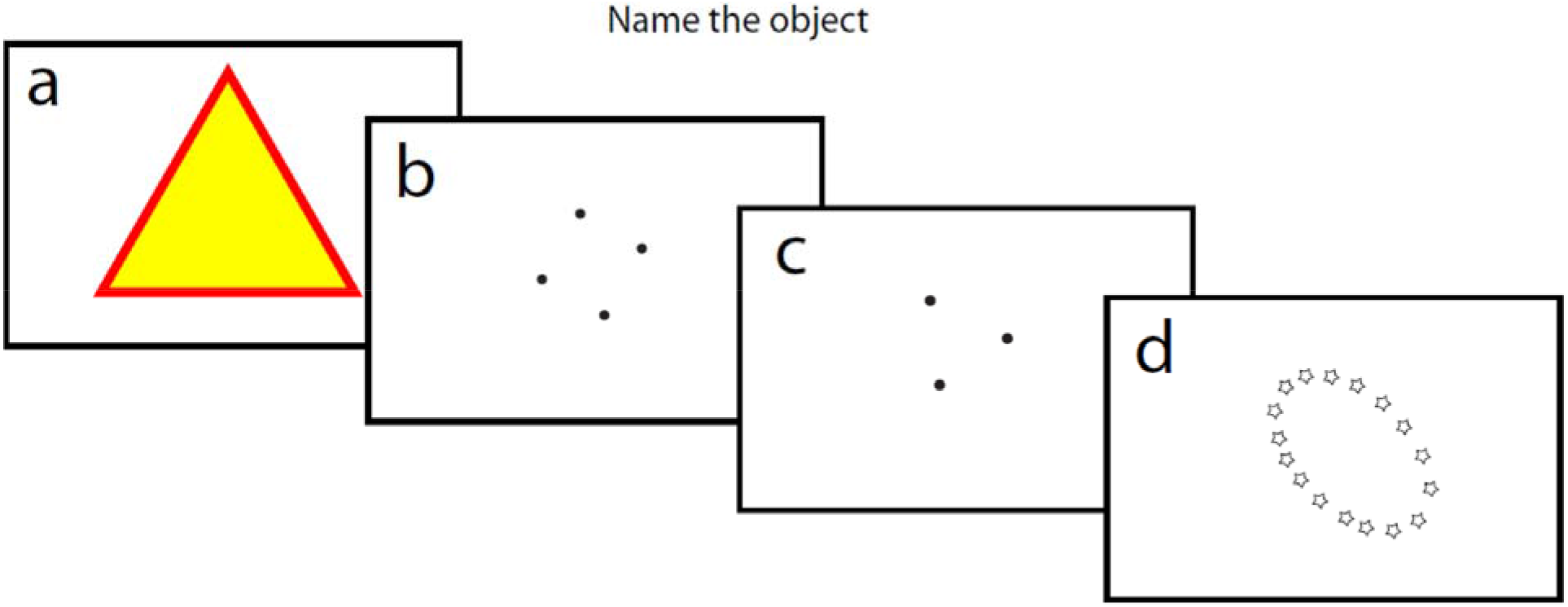
A thought experiment where the subject is asked to name the object (triangle, square, or ellipse) while ignoring the nature of the elements making up the individual objects.

#### Category uniqueness and invariance

Each object ensemble is assumed to be tuned to a particular category so that its population responds vigorously and similarly to all, say, chairs, regardless of shape, size, orientation or texture, while responding weakly, if at all, to all other objects. To go back to our experiment, are we to assume that the subject can name the various objects because the triangle, square and ellipse evoke activity in three corresponding ensembles, and that the “triangle ensemble” is responding similarly to the two greatly different triangles (Fig. 6, a, c) but does not respond to the square (Fig. 6b) despite its similarity to the triangle in Fig. 6c?

To expand on the latter point, let’s look at Fig. 7. There, for each triangle property such as size or orientation, a number of possible variations are displayed. If we, conservatively, consider 15 variations of 10 properties, we end up with 15 possible triangles. Thus the “triangle ensemble” is capable not only of responding effectively to triangles while ignoring many thousands of other objects, but also of generating a stereotypic triangle-categorizing response to countless possible appearances of triangles (Fig. 7).

**Figure 7.**
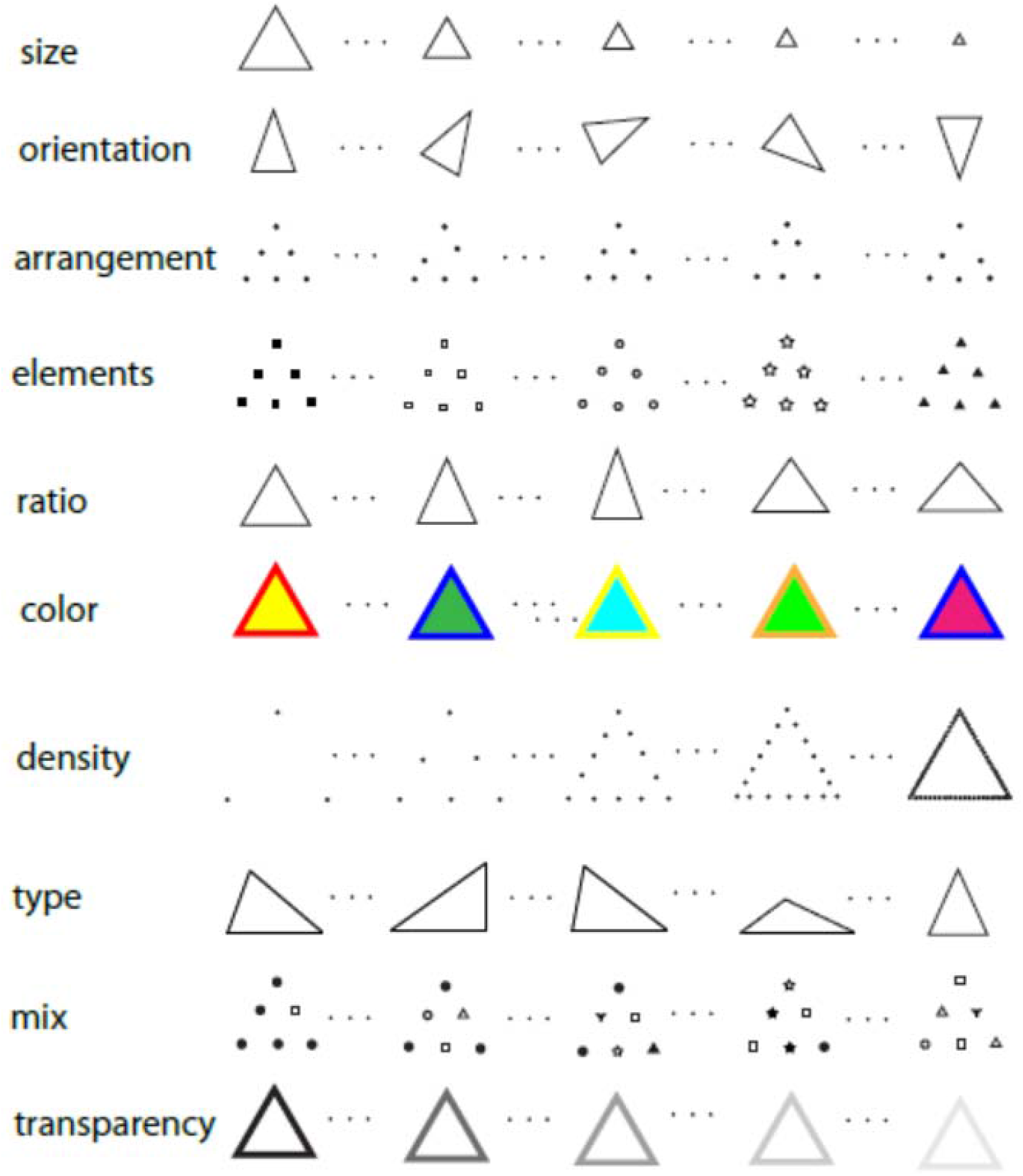
A demonstration of the almost unlimited variants that are clearly identified as triangles.

#### Within-category uniqueness and invariance

Even more challenging is the contradictory requirements of an “object-ensemble” to generate a category identifying response to all individual members of the category and at the same time maintain each exemplar’s individual identity. This goes beyond unlikely; it is impossible.

In the above example (Fig. 7), the “triangle ensemble” is assumed to generate a limited set of responses that distinguish all 10^15^ variants from any other object and at the same time produce 10 different responses to account for our ability to discriminate between all exemplars. Not only impossible but absurdly so.

#### Local changes

In addition to the above requirements objects are recognized and discriminated from other objects not only when undergoing global changes in, say, size or orientation, but also when there are local changes. Let us consider another simple thought experiment (Fig. 8) where the subject is asked to determine whether the two faces, one with glasses and one without, represent the same person. According to the ECE hypothesis, the face-ensemble responds uniquely to any of the vast numbers of faces that we can discriminate between. However, faces can change many local details (hair style, glasses, makeup, or facial expressions) yet maintain individual identity. So now, the face-ensemble must give the same individual-related response and ignore not only countless global variations but countless local variations as well. It is clear that such a performance by an ECE is impossible. Which brings us to a fundamental failure of the ECE approach.

### The ECE hypothesis by its very nature cannot explain our visual perception

Present results and the analysis of the above thought experiments, demonstrate that the expert cell ensemble hypothesis fails to account for our ability to parcel objects into many thousands of categories, and to distinguish an object that can appear in infinite global and/or local variations. Next, I argue that the very idea of a convergent, feature-extracting, code-generating system contradicts our perceptual abilities.

Our perception of spatial elements, from the tiniest speckles to complex objects is, by and large, parallel in space, simultaneous, comprehensive, and veridical. We perceive correctly the position of, say, any two elements within visual space as well as their relative size, luminance, orientation, or color. Without the ability to simultaneously perceive many elements, no complex structure such as a face could have been perceived as an integrated holistic object. When we see an object we perceive an integrated entity and at the same time we can see, with great accuracy, all the elements that make up that object. In fact, it is this very ability of visualizing the smallest constituent elements of any object that allows us to discriminate and recognize individual ones. These self-evident and elementary observations are, nonetheless, in stark contradiction to the mechanisms and predictions of the ECE hypothesis. Next, let us contrast the ECE main tenets (bold headings) with perceptual evidence.

#### A necessary mechanism for object representation and perception is feature extraction via convergence and integration

However, we do not see the world as a collection of lines and edges as a V1 output-based perception would predict. Moreover, if, as practically all investigators believe, our perception is based on IT cell responses, we would know, at best, that a certain object is before us, but we would not see an actual object with all of its constituent elements. In other words, unlike the feature-extracting prediction, we always perceive both the forest and the trees.

#### Object representation and perception are based on sparse encoding by small cell ensembles in the IT cortex

According to the ECE hypothesis an object is uniquely encoded by the collective neural activity of a few hundred cells in the IT cortex. Put differently, the expert ensemble responses represents an input, a spatially distributed light intensities―an object― by the firing patterns of some IT cells where all spatial, contrast and wavelength information is irretrievably lost.

In visual areas V1-V4, RFs are organized retinotopically so that although many details are integrated over by hierarchical convergence, the rough spatial outlines of represented objects are maintained. At the top of the hierarchy, the IT cortex, there is a fundamental change; RFs are no longer retinotopic but cover a large portion of the visual field. This means that “expert” IT cells response patterns must be interpreted purely as a code. This total break from retinotopic representation and from any spatial information contradicts our perception where spatial locations, organization, and details are clearly evident.

#### The activity pattern of expert cell ensembles is invariant to global and many local changes in the object’s appearance

One of the essential requirements of the ECE hypothesis and, in fact, of any computational object recognition system, is that object representation be invariant to the many possible global changes in the object appearance such as size, orientation, or luminance. As argued above, given the huge number of possible global and local variations it is impossible for any computational system to generate the same response for all possible variations of a given object. More importantly, we perceive all and every variation in any object that we see. As demonstrated in the present experiments and in our daily experience an object is recognized when its image is large or small, tilted or not, brightly illuminated or not.

#### To summarize: what makes visual perception unique, powerful, and infinitely flexible, is what fails all feature extracting/computational/encoding systems

We are able to perceive, discriminate, and recognize a very large number of different object classes, as well as objects within a class where each may appear in countless variations. While computational systems struggle to eliminate variability and use extensive learning to generate dedicated units so that any object is forced into a canonical representation―objects out there in the world are perceived, by and large, as they are; relative sizes, positions, orientations, and luminance are accurately preserved. Admittedly, there are many minor exceptions to this rule such as the inverted retinal image, imaginary borders, interactive effects and more, but the basic structure of the objects is veridically preserved. So while computational models try to extract from the image its invariant essential features, we perceive objects in their raw format; all details, not just “characteristic features”, are there, all variants are there, all shades of grey or any combination of colors are perceived―all of which not only do not “mask” the object but are essential to its recognition. In short, all the crucial features in computational models―convergence, feature extraction, invariance, encoding―are not found in our conscious visual perception, while what makes our perception so successful―detail preservation, discrimination and recognition of huge number of objects that can appear in countless variations―are the characteristics that all models fail to account for.

#### If object representation and recognition are not achieved via a convergent, feature-extracting, encoding, computational system, how is it done?

Let me stress that demonstrating the utter inability of biologically-plausible models, such as ECE, as well as any computational, encoding model, to explain our perceptual abilities is by itself highly important. Using computer-friendly models that ignore the obvious abilities of visual perception has swayed the field into a stagnating, dead-end state. By realizing the futility of computational modeling we are free to look for alternative explanations―ones that would be driven by perceptual evidence rather than by the tools at hand. To paraphrase “Maslow’s hammer” adage, if you have a computer, every perceptual phenomenon is computable. So here are a few thoughts in a search of an alternative explanation.

### Toward an alternative explanation

#### Object perception: parallel, instantaneous and simultaneous, detailed and holistic

##### Parallel

The parallel spatial elements making up an object are projected on the retina and this parallel representation is kept largely intact in primary visual cortex. As information flows downstream from V1 through cortical areas V2 and V4, spatial resolution degrades and is completely lost in the IT cortex. Our perception of spatial objects is parallel as is V1 representation; we do not perceive just edges or borders that V1 output is assumed to represent, or the rough low spatial frequency output of V2 and V4, and certainly not a coded representation of objects attributed to the IT cortex.

##### Instantaneous and simultaneous

There are many studies showing that humans are able to recognize or discriminate briefly flashed (6.25-20ms), fairly complex stimuli displaying animals (Thorpe et al., 1996; Faber-Thorpe et al., 2001) or faces (Rolls et al., 1994; Keysers et al., 2001). The latter showed comparable results for single cell responses and human performance. Recent studies by Greene and co-workers (Greene and Ogden, 2012; Greene and Visani, 2015) showed that human subjects can easily discriminate various shapes such as letters or animals that are flashed for extemely short (50 *µsec* or less) durations. Even using masks with a 40*ms* stimulus onset asynchrony does not prevent face discrimination or identification at >90% success (Purcell and Stewart, 1988; Rolls et al., 1994; Faber-Thorpe et al., 2001; Or and Wilson, 2010). Of particular note is my recent work (Gur, 2018) where very small faces (<19’) were successfully discriminated when presented for a mere 17ms. In addition, we have recently shown that Sloan characters could be recognized at a 1’ acuity when presented for 0.5ms (Rosen et al., 2020). Present results where 0.6-2.4º faces presented for 83ms (followed by a 100ms mask) were recognized at >90% success are another demonstration of our ability to perceive and process briefly displayed complex objects. Our capacity to relate spatial information presented for such short times at disparate visual field loci means that perceptual interactions and comparisons between object elements is simultaneous and, practically, instantaneous.

##### Detailed and holistic

These two characteristics which to any integrative/computational system are contradictory since the whole is derived by integrating over its parts, are very much the way we perceive the world. It is the visual system ability to perceive space simultaneously and in parallel that leads to the holistic yet detailed capacity; the world is perceived almost as is in a flash; all the elements as well as their relative features (size, shape, orientation, brightness, etc.) are available to the system for an immediate decision―a triangle or a square? A building or a face? Such a discrimination between objects or such a retention of details would not have been possible if minute spatial data were integrated and serially sent across many synapses for tens or hundreds of milliseconds to areas downstream from V1.

For a first-hand experience of those perceptual abilities the reader is encouraged to look at Demos1-3 in the supplemental file (Demos.docx).

#### What might be the neural basis of our unique perceptual abilities?

Responses in primary visual cortex to images falling on the retina can account for parallel, instantaneous, and detailed perception. An image generates, instantaneously, a corresponding parallel, spatial pattern in V1 where all details, spatial relations, luminance levels, colors and more are preserved. Since V1 is the only cortical area where RF size is compatible with our spatial resolution, its responses must be the neural basis of our visual perception (for a detailed discussion of this issue see Gur, 2015).

### If V1 input patterns are the basis of visual perception, what is it the rest of the visual cortex doing?

While the representation of the 2D visual image in V1 is the basis of our detailed, accurate, parallel perception, there is much more than that to our perception. What our brain adds to the 2D image is scene understanding and interpretation. It is necessary to recreate the 3D spatial information that has been reduced to a 2D one. This is done mainly by making use of monocular cues for depth perception―cues such as perspective, texture, absolute and relative distance and size, motion, and more. We use our understanding of the world to realize that an incomplete object is an actual one that is partially hidden by another object, and we utilize our memory to relate an acute image to a stored one, or to a concept (a triangle). Clearly, for making sense of the flat V1 representation a great deal of processing and neural specialization is required. Indeed, for that processing, the integrative feature-extracting properties of single cells, from V1 simple cells onward are essential. Put differently, to perceive that there is an object on our left moving slowly toward us, and that this object is a small human being, we rely on global information gleaned from responses of cells located, for example, in areas V2, V4 (size, orientation, location), V5 (motion, direction), and IT (human). To perceive that this object is our son Jacob, we must rely on V1 pre-integration input patterns.

### A parallel, simultaneous neural activity directly generates a parallel, simultaneous conscious perception

Given the above considerations, we can now outline the neural elements that generate object perception and recognition. In V1, neurons responding to the various elements that make up an object are activated simultaneously and in parallel. After a short delay, many neurons in various visual areas respond to more integrative holistic features such as size, orientation, or motion. This matrix of active cells is the substance of our conscious visual perception.

#### We cannot bypass the mystery of consciousness

A popular notion is the distinction between the “hard” and the “easy” problems of consciousness (Chalmers, 1995). Whereas the hard problem relates to the very experience of consciousness, the easy one are phenomena such as sensory perception that, in principle, can be explained by computational or neuronal mechanisms. As shown above, conscious visual perception cannot be explained by such mechanisms. Rather, it is likely that perceptual abilities that are evident to the conscious viewer emerge from the same mechanisms that generate conscious experience *per se*. Both manifest the unexplained transformation from objective brain activity to subjective, private conscious perception and experience. The pattern of responding visual cells corresponds only to our ability to perceive localized, point-wise details but not to our perception of holistic information that is shaped and determined by details (fifty dots that can look like a rabbit or a turtle), or to our ability to relate and compare local information across the visual field. Such abilities cannot be explained by convergence to omnipotent “object cells” or by local interactions carried by axons connecting each locus in the VF to all other loci. The discrepancy between objective neural activity and subjective perception is so striking that we should accept the mysterious transformation from objective neural responses to subjective perception, accept that we do not understand how it is done, and not try to avoid the issue by relying on mechanisms that are non-biological, and do not account for our perceptual performance.

#### Perception exhibits field-like properties

Since the unique properties of conscious perception cannot be explained by lateral, feed-forward, or feed-back axonal transmission, it may be useful to think in terms of a “wireless” mechanism―a field. An electric field, for example, is shaped by the number and strength of its electrical charges; changes in the charges’ characteristics instantly (at the speed of light) affect the shape of the force distribution of the whole field, yet the properties of individual charges (location and strength) are preserved. We don’t know what sort of “neural field” we experience when we perceive what is out there but to better understand it we have to account for the unique properties of conscious visual perception rather than rely on well-understood, readily simulated mechanisms that ignore perceptual reality. Let me add that suggesting some sort of a field to explain conscious phenomena is not new (cf. Libet’s, 2006, “conscious mental field”) and is usually based on introspection about the nature of conscious experience. Pockett (2017) hypothesized that consciousness is identical with spatially extensive electromagnetic fields which require, *inter alia*, feed-back to primary sensory areas. However, both spatial extent and feed-back are not consistent with our ability to consciously perceive small individual elements within a time frame that is too short for any top-down feed-back. Here, the notion of a “neural field” is proposed as arising naturally, almost by default from empirical evidence and functional anatomy reality showing that mechanisms based on axonal transmissions and interactions fail to account for conscious visual perception.

## Supporting information

Supplemental Demos

