## Supplemental Demos for "Psychophysical evidence and perceptual observations show that object recognition is not hierarchical but is a parallel, simultaneous, egalitarian, non-computational system"

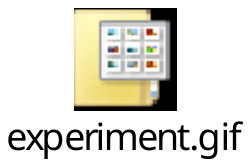

A demo showing experimental procedure; here the stimulus is presented for a 100ms (somewhat longer than the 67ms used in the experiments) followed by a 100ms mask and a 400ms pause. Click to activate. Demos presentation times are based on a 60 Hz refresh rate.

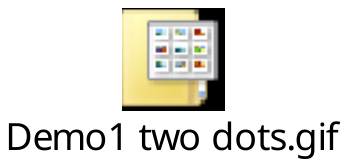

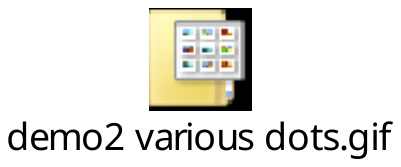

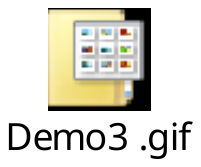

Three demos showing our ability to integrate across space briefly presented elements. In demo1 we can easily see two dots of unequal size that change position every 17ms. In demo2 each frame is displayed for 34ms and we see the change in the dots’ number, size, color, orientation, and location. In demo3, four identical and one outstanding faces are displayed for 34ms each with the whole series repeating every 50ms. The outstanding face can be readily observed.
